# GrapHi-C: Graph-based visualization of Hi-C Datasets

**DOI:** 10.1101/156679

**Authors:** Kimberly MacKay, Anthony Kusalik, Christopher H. Eskiw

## Abstract

**Background:** Hi-C is a proximity-based ligation reaction used to detect regions of the genome that are close in 3D space (or “interacting”). Typically, results from Hi-C experiments (whole-genome contact maps) are visualized as heatmaps or Circos plots. While informative, these visualizations do not intuitively represent the complex organization and folding of the genome in 3D space, making the interpretation of the underlying 3D genomic organization difficult. Our objective was to utilize existing tools to generate a graph-based representation of a whole-genome contact map that leads to a more intuitive visualization.

**Methodology:** Whole-genome contact maps were converted into graphs where each vertex represented a genomic region and each edge represented a detected or known interaction between two vertices. Three types of interactions were represented in the graph: linear, intra-chromosomal (*cis*-), and inter-chromosomal (*trans*-) interactions. Each edge had an associated weight related to the linear distance (Hi-C experimental resolution) or the associated interaction frequency from the contact map. Graphs were generated based on this representation scheme for whole-genome contact maps from a fission yeast dataset where yeast mutants were used to identify specific principles influencing genome organization (GEO accession: GSE56849). Graphs were visualized in Cytoscape with an edge-weighted spring embedded layout where vertices and linear interaction edges were coloured according to their corresponding chromosome.

**Results:** The graph-based visualizations (compared to the equivalent heatmaps) more intuitively represented the effects of the *rad21* mutant on genome organization. Specifically, the graph based visualizations clearly highlighted the loss of structural globules and a greater intermingling of chromosomes in the mutant strain when compared to the wild-type. The graph-based representation and visualization protocol developed here will aid in understanding the complex organization and folding of the genome.

## 1 Introduction

One of the major problems in the genomic era is understanding how genomes are organized and folded within cells. The organization of the genome, specifically the close physical proximity of genetic elements located distally on the same or different chromosomes, greatly impacts cellular processes such as development and disease progression. Knowledge of what interactions are occurring and how they are mediated is essential to understanding various genome functions such as the regulation of gene expression. The biological experiment Hi-C [2,6] (or one of its derivatives [3-5,12]) can be used to detect regions of the genome in close 3D spatial proximity. This close physical proximity is often referred to as an “interaction” between two genomic regions. Hi-C is able to detect interactions between regions of the genome on the same chromosome (intra-chromosomal or *cis*-interactions) as well as interactions between regions of the genome on different chromosomes (inter-chromosomal or *trans*-interactions). Briefly, Hi-C involves chemically cross-linking regions of the genome that are in close physical proximity. Restriction enzyme digestion and ligation is then preformed on the cross-linked regions to generate novel chromatin/DNA complexes which can be identified by next generation sequencing. The resultant sequence reads are mapped to a reference genome [1] to generate a whole-genome contact map which represents all of the detected Hi-C mediated interactions.

The results of a Hi-C experiment are often encoded as a symmetric *N* × *N* matrix (referred to as a whole-genome contact map) where N is the number of genomic “bins” that the genome is partitioned into. Each genomic bin represents a linear region of genomic DNA based on the experimental resolution. For instance, a Hi-C experiment in fission yeast that is able to attain 10 kB resolution will generate 1258 genomic bins, where each bin represents roughly 10 kB of linear DNA sequence. In general, the number of genomic bins is approximately equal to the total genome size divided by the Hi-C experimental resolution. Each cell (*CM_i,j_*) of the whole-genome contact map records how many times the genomic bin *i* was found to interact with the genomic bin *j*. This is often referred to as the frequency of the interaction between the two bins.

Typically, whole-genome contact maps are visualized as heatmaps (Figure 2A, B) or Circos plots [11]. While informative, these visualizations do not intuitively represent the complex organization and folding of the genome in 3D space. This makes it difficult to quickly understand the underlying 3D genome organization represented by the contact map. Our hypothesis was that representing and visualizing whole-genome contact maps as a graph would lead to a more intuitive visualization of Hi-C data when compared to typical methods. We developed a package called GrapHi-C (pronounced as “graphic”) for visualizing Hi-C data as a graph. GrapHi-C utilizes a graph-based representation of a whole-genome contact map and existing tools to more intuitively visualize Hi-C data. Briefly, whole-genome contact maps were converted into a graph where each vertex represented a genomic region and each edge represented an interaction between two vertices. In addition to edges representing the detected *cis*-and *trans-* interactions, edges representing the linear connections between genomic regions were also included. Since these linear edges represent *bonafide in vivo* connections between the genomic bins, this adds an additional biological constraint to the graph-based visualization. Overall, the graph-based visualization protocol developed here was able to intuitively represent the complex genomic organization for a given whole-genome contact map.

**Fig.2.**
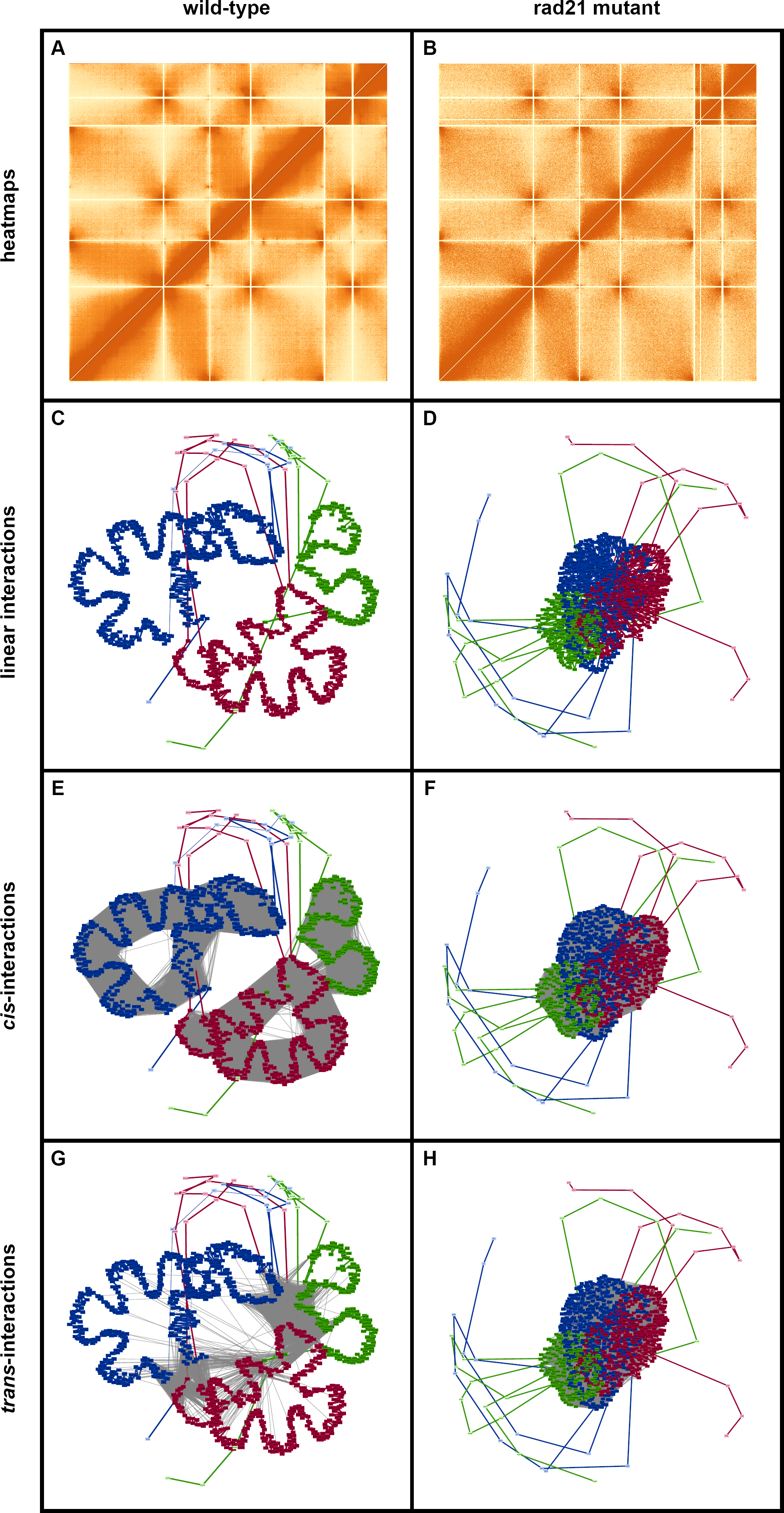
Comparison of Heatmaps and Graph-Based Visualization for Wild-Type and *rad21* Mutant Fission Yeast Whole-Genome Contact Maps. Visualizations of the whole-genome contact maps for the fission yeast wild-type and *rad21* mutant are displayed in the left and right columns, respectively. Panels A and B are the heatmaps generated with ggplot2 in R that correspond the the contact maps. Panels C-H are the graph-based visualizations where vertices and linear interactions were coloured according to their corresponding chromosome. The grey dashed lines represent cis-interactions (Panels E and F) and trans-interactions (Panels G and H). Graphs were visualized in Cytoscape using a edge-weighted spring embedded layout.

## 2 Methods

### 2.1 Graph-Based Representation

Building on a previously proposed graph-based representation of a whole-genome contact map [11], each vertex in the graph represents a genomic region (bin) and each edge represents a detected or known interaction between two bins. Specifically, edges representing linear, *cis-* and *trans-* interactions were depicted in the graph. Each edge was weighted with the experimental resolution (for linear interactions) or the associated interaction frequency (for *cis-* and transinteractions). A formal description of the developed graph based representation is presented below in Formal Model 1.

##### Formal Model 1

Formal description of the graphical representation of a whole-genome contact map

##### Whole-Genome Contact Map Definitions

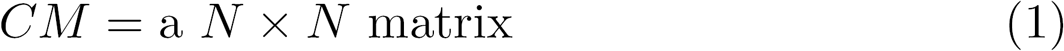

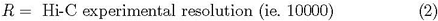

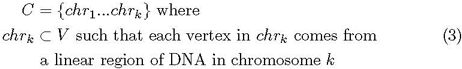

##### Graph Definitions

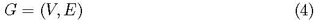

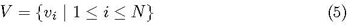

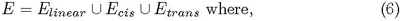

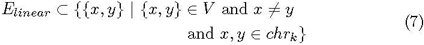

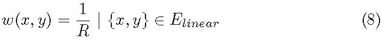

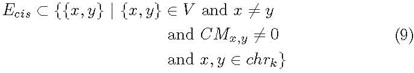

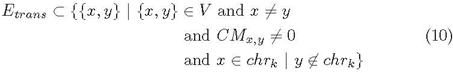

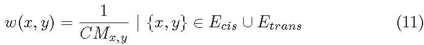

#### 2.2 Visualization Protocol

A Perl script was developed that is able to convert a normalized whole-genome contact map into an adjacency matrix based on the graph representation described above (available at: https://github.com/kimmackay/GrapHi-C). The output of this script can then be input into Cytoscape [8] and visualized as a graph. The *S. pombe* whole-genome contact maps were visualized in Cytoscape using an edge-weighted spring embedded layout with the default parameters. A general overview of the developed protocol is depicted in Figure 1.

**Fig.1.**
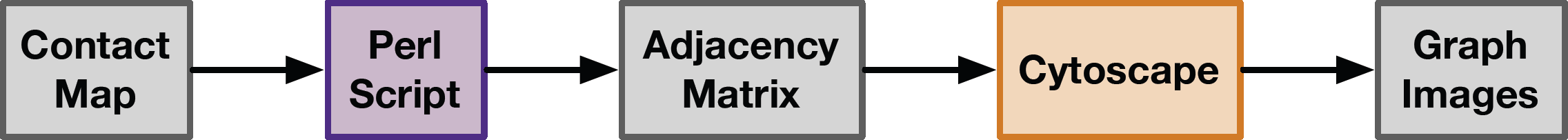
Overview of the protocol for visualizing whole-genome contact maps as a graph. Each step of the workflow is indicated in a box where the different colours correspond to: data input and output (grey), developed Perl script (purple), and an existing tool (orange).

### 3 Results & Discussion

To demonstrate the usability of the developed graph-based representation and protocol, we used it to visualize whole-genome contact maps from an existing fission yeast dataset. Normalized *S. pombe* whole-genome contact maps were downloaded from the Gene Expression Omnibus database (accession number: GSE56849) [7]. Specifically, we utilized the 999a wild-type and the *rad21* deletion contact maps. An adjacency matrix for each contact map was generated using the Perl script described above. The resultant graphs were visualized in Cytoscape with an edge-weighted spring embedded layout (Figure 2C-H). Vertices and the linear edges were coloured according to their corresponding chromosome. Vertices along the periphery of the graph images correspond to genomic bins that represent centromere and telomere regions. Since these regions are highly condensed and repetitive (making the DNA difficult to access and map) no interaction data was reported for them. For the purpose of comparison, the heatmaps for the 999a wild-type and the *rad21* deletion contact maps were generated using ggplot2 [10] in R [9] (Figure 2A,B). These heatmaps represent the standard, existing approach for the visualization of Hi-C data.

The heatmaps that correspond to the wild-type and *rad21* mutant whole-genome contact maps clearly demonstrate how this traditional form of visualization does not intuitively represent the complex organization and folding of the genome in 3D space (Figure 2A, B). This makes it challenging to generate hypotheses about how the differences in the wild-type and mutant contact maps could impact genome organization. On the contrary, the graph-based visualizations (Figure 2C-H) clearly depict the differences in the wild-type and mutant whole-genome contact maps. Furthermore, as discussed below the graph-based visualizations more intuitively represent how these distinct contact maps may affect genome organization. It should be noted that for the graph-based visualizations, all of the edges (that correspond to *cis*-, *trans-* and linear interactions) were used to generate the graphs in each panel. For the purpose of visualization, each panel highlights only one type of edge while the other two are hidden. The resultant visualizations for the *rad21* mutant strain (Figure 2D, F, H) appear to be very similar since the mutant contact map has smaller interaction frequency values (as compared to the wild type) which results in the nodes being placed closer together in the edge-weighted spring embedded layout. This density makes it difficult to see the differences when the edges corresponding to the *cis*-, *trans-* or linear interactions are highlighted.

Overall, comparison of the graph-based visualizations for the wild-type and *rad21* mutant whole-genome contact maps highlights the loss of structural globules (intra-chromosomal structures) and a greater intermingling of chromosomes in the mutant strain [7]. Specifically, the graph-based visualizations (compared to the equivalent heatmaps) made it easier to quickly identify the principles of genome organization and associated biological effects of the *rad21* yeast mutant that were discovered in the original study. This suggests that the developed graph-based representation and visualization is a biologically valid way to represent whole-genome contact maps.

### 4 Conclusion

In this manuscript, we built on previously proposed graph-based representations of whole-genome contact maps and described a novel protocol for visualizing Hi-C data as a graph. In addition to edges that represent the detected cis-and *trans-* interactions, we chose to include edges between each sequential genomic bin within a chromosome to better represent the linear extent of the genome. This adds additional, verified in *vivo* biological constraints to the visualization. We developed a simple Perl script that can be used to convert a whole-genome contact map into an adjacency matrix related to the developed graph-based representation. This matrix can then be input into Cytoscape (an existing tool) for visualization.

Overall, the developed graph-based visualization of the whole-genome contact maps (compared to the equivalent heatmaps) made it easier to quickly identify the changes in genome organization seen in the yeast mutants identified in the original study. Future work will focus on extending this visualization to allow for the vertices to be coloured according to complementary-omics datasets (such as gene expression) and produce a 3D graph-based visualization. Additionally, we will apply it to organisms with larger genomes to see how well this visualization scales for larger whole-genome contact maps. Not only does the graph-based representation of Hi-C data lead to a more intuitive visualization, it also has the potential to lead to new ways of analyzing whole-genome contact maps by leveraging the extensive amount of graph theoretic literature.

## Key Points

— A formal graphical representation was developed for a whole-genome contact map. Unlike previously proposed graph-based representations, this formal model includes linear interactions between sequential genomic bins within a chromosome.
— A Perl script was developed that takes a whole-genome contact map as input and outputs an adjacency matrix corresponding to the formal graph model described above.
— A new protocol was developed that utilizes Cytoscape to visualize whole-genome contact maps as a graph.
— The developed graph-based visualization of the whole-genome contact maps (compared to the equivalent heatmaps) made it easier to intuitively understand the underlying principles of 3D genome organizations that were identified in a previous study.

## Data and Software Availability

Datasets analyzed in this study are available in the Gene Expression Omnibus database (accession number: GSM1379427;https://www.ncbi.nlm.nih.gov/geo/query/acc.cgi?acc=GSM1379427). The Perl script used for converting a whole-genome contact map into an adjacency matrix (based on the graph representation described in this manuscript) is available at https://github.com/kimmackay/GrapHi-C/. This work is licensed under the Creative Commons Attribution-NonCommercial-ShareAlike 3.0 Unported License. To view a copy of this license, visit http://creativecommons.org/licenses/by-nc-sa/3.0/ or send a letter to Creative Commons, PO Box 1866, Mountain View, CA 94042, USA.

## About the Authors

Kimberly MacKay is a Ph.D. student at the University of Saskatchewan in the Department of Computer Science. Christopher H. Eskiw is an Assistant Professor at the University of Saskatchewan in the Department of Food and Biproduct Sciences. Anthony Kusalik is a Professor in the Department of Computer Science, a member of the Division of Biomedical Engineering and the Director of the Bioinformatics Program at the University of Saskatchewan.

## Acknowledgements

Not Applicable.

